# Tet3 Mediates O-GlcNAcylation-Induced Remyelination

**DOI:** 10.1101/2025.01.24.634702

**Authors:** Yawen Li, Pingping Li, Yixuan Song, Mingliang Zhang

## Abstract

The failure of oligodendrocyte precursor cells (OPCs) to generate mature oligodendrocytes represents a major hurdle impeding myelin repair. However, the cellular determinants for OPC differentiation and the pharmacological targets driving remyelination remain rudimentary. Here, we demonstrate that O-GlcNAcylation intrinsically drives remyelination, in which Tet3 is one downstream effector. Enhanced O-GlcNAcylation significantly promotes OPC differentiation, whereas conditional depletion of Ogt in OPCs seriously impairs myelin formation in the developing central nervous system (CNS). Intriguingly, O-GlcNAcylation in OPCs positively correlates with the myelin status, and enhanced O-GlcNAcylation dramatically accelerates remyelination. Proteomic analyses reveal a cohort of proteins uniquely O-GlcNAcylated in OPCs. Among these, Tet3 is exclusively O-GlcNAcylated in OPCs within CNS, which in turn modulates its demethylation activity, thereby regulating myelin gene expression and remyelination. Our findings highlight the crucial role of the Ogt-Tet3 axis in OPC differentiation and suggest O-GlcNAcylation as a druggable target for treating demyelinating diseases.

## INTRODUCTION

In the central nervous system (CNS), the principal role of oligodendrocyte precursor cells (OPCs) is to respond to external cues and produce myelinating oligodendrocytes (OLs) (*1,2*). Mature OLs then ensheathe the axons of neurons with myelin (*3*). By doing that, OPCs enable fast saltatory impulse propagation, orchestrate nearby neuronal activity, and modulate brain function (*4*). Many conditions (e.g. aging, autoimmune, etc) could destroy myelin and ultimately lead to demyelinating diseases (*5–10*). Loss of myelin would not only disrupt the axonal energy metabolism but has been recently considered a potent driver for neurodegenerative disorders (*11*). In addition, excessive myelin degradation is a major change observed in aged non-human primate brains and would result in deleterious effects on neighboring neurons, which may eventually trigger brain aging (*12*). Therefore, maintaining and rejuvenating the intact myelin is indispensably important to brain function (*13*).

In the conditions of demyelination, OPCs regenerate myelin and recover the function of neurons and the brain (*6,14*). Although myelin could be partially restored, the process of remyelination is inefficient (*9*). Clinical studies revealed that OPCs are abundant in regions of demyelination, yet fail to differentiate (*15*). Thus, failure to regenerate myelin by OPCs represents a major hurdle impeding myelin repair (*15,16*). The current medications for treating demyelinating diseases, however, are predominantly immune-modulating (*1,17*). Thus, to develop the medications promoting myelin repair by directly targeting oligodendroglia is highly desired. Unfortunately, the intrinsic determinants modulating OPC differentiation and the pharmaceutical targets for clinical intervention remain largely elusive.

O-GlcNAcylation is an abundant and dynamic post-translational modification (PTM) enriched in the brain (*18–20*). O-GlcNAcylation is catalyzed by a single pair of enzymes, O-GlcNAc transferase (Ogt) which transfers β-N-acetylglucosamine (O-GlcNAc) to the hydroxyl groups of serine/threonine residues of nucleocytoplasmic proteins and O-GlcNAcase (Oga) that is responsible for removing this modification (*21*). O-GlcNAcylation plays a pivotal role in neurodevelopment and rejuvenation (*22–26*).

Mechanistically, O-GlcNAcylation modulates the stability, locations, and/or catalytic activity of its modified proteins, which, in turn, regulates diverse biological processes (*18,27*). Given the downstream targets and biological consequences are context-dependent, how O-GlcNAcylation works in a certain type of cell over a specific biological process, such as the differentiation in OPCs, is largely unclear.

Here, we identify O-GlcNAcylation as an intrinsic driver promoting remyelination. O-GlcNAcylation increases dramatically in OPCs after birth, when robust myelination is initiated. Conditional knockout of *Ogt* in OPCs led to reduced O-GlcNAcylation, impaired OPC differentiation, and consequently attenuated myelin formation in developing CNS. The extent of O-GlcNAcylation in OPCs is positively correlated with myelin status. Upregulation of O-GlcNAcylation dramatically promoted OPC differentiation in culture and induced remyelination in a mouse model of demyelination. The initial mechanistic study reveals that O-GlcNAcylation orchestrates repertories of the pro-myelinating program through its modified proteins. Among these, Tet3 is verified experimentally as one downstream effector of Ogt. Intriguingly, O-GlcNAcylation affects the demethylation activity of Tet3, which in turn regulates the transcription of myelin genes and consequently remyelination. Collectively, our finding not only identifies O-GlcNAcylation as a cellular determinant of OPCs for remyelination but also reveals the mechanistic insight of Tet3 on myelin formation. Importantly, our discovery highlights the potential of targeting O-GlcNAcylation as an effective treatment for demyelinating diseases.

## RESULTS

### O-GlcNAcylation Modification for Proteins in OPCs

To explore the function of O-GlcNAcylation on myelin formation, we first assessed the expression of Ogt and O-GlcNAcylation in OPCs. We established a step-wise differentiation assay of mouse embryonic stem cells (ESCs) towards oligodendroglial cells. By immunostaining, O-GlcNAcylation (evidenced by RL2-reacting proteins) was detected in ESCs and neural progenitor cells (NPCs) (Fig. 1A), as previously reported (*23,28*). We observed a high percentage of OPCs express Ogt or are positive for RL2 (Fig. 1A and fig. S1A). *Ogt* expression in OPCs was further confirmed by quantitative real-time PCR (qRT-PCR) analysis (fig. S1B). By Western blotting, we detected the O-GlcNAcylation, as well as Ogt expression, in OPCs (Fig. 1B and fig. S1, C and D). These results demonstrated a high abundance of O-GlcNAcylated proteins in OPCs in culture.

**Fig. 1.**
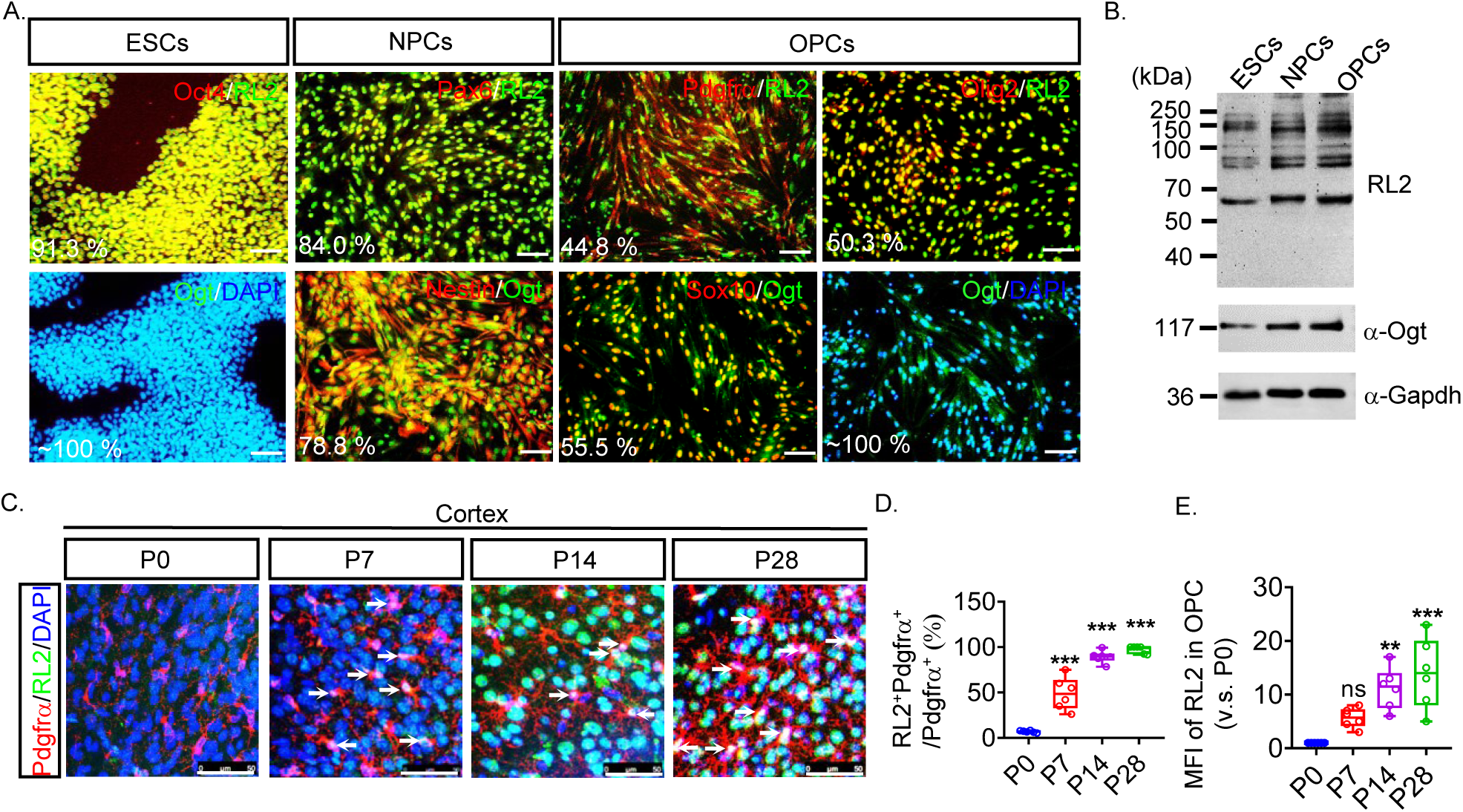
O-GlcNAcylation Modification for Proteins in OPCs. (**A**) Immunostaining analysis for Ogt or the O-GlcNAcylated proteins in ESCs, NPCs, and OPCs. The percentage of cells co-expressing indicated cell type-specific markers (Oct4 for ESCs, Pax6, and Nestin for NPCs, and Pdgfrα, Sox10, Olig2 for OPCs, respectively) and RL2 (anti-O-GlcNAcylation) or Ogt was shown in (**A**). Scale bars, 50 μm. n=6 independent biological experiments. (**B**) Western blot analysis for O-GlcNAcylated proteins (upper panel), Ogt (middle panel), and Gapdh (bottom panel) in indicated cell types. n=5 independent biological experiments. (**C-E**) Immunofluorescence analyses show the O-GlcNAcylation modification in Pdgfrα^+^ OPCs from the cortex of mice at P0, P7, P14, and P28, respectively (**C**). Scale bars, 50 μm in (**C**). The percentage of RL2^+^ OPCs (RL2^+^Pdgfrα^+^/Pdgfrα^+^ cells) was quantified in (**D**). The fold enrichment of the mean fluorescence intensity (MFI) of RL2 in Pdgfrα^+^ OPCs was quantified and compared to that in P0 (**E**). n=6 mice for each time point. All the quantification data are presented as mean ± SEM. *p*-values were calculated using one-way ANOVA test (**D, E**). \**p* < 0.05, \*\**p* < 0.01, \*\*\**p* < 0.001, ns: not significant.

To assess O-GlcNAcylation in OPCs *in vivo*, we analyzed the OPCs in the cortex at postnatal day 0 (P0), P7, P14, and P28, a time course covering robust myelin development after birth (Fig. 1, C to E). At P0, we hardly detected the O-GlcNAcylated proteins in the platelet-derived growth factor receptor alpha (Pdgfrα)-expressing OPCs (Fig. 1, C to E). Myelination is initiated at about P7 in CNS, and concomitantly the O-GlcNAcylation became evident in OPCs (Fig. 1, C to E). Meanwhile, we observed a significant augment of O-GlcNAcylation in OPCs from P7 to P14 (Fig. 1, C to E), when robust myelination started, implying an involvement of O-GlcNAcylation in myelin development. We detected the increasing expression of RL2 in OPCs, which was also observed in OPCs in the corpus callosum (CC) (fig. S1, E to G). Together, these data demonstrate that Ogt O-GlcNAcylates proteins in OPCs *in vivo*.

### O-GlcNAcylation Regulates OPC Differentiation *in vitro*

To understand the role of O-GlcNAcylation, we assessed its function on OPC differentiation. We collected the single ESC colony-derived OPCs with doxycycline (Dox)-inducible expression of CRISPR-Cas9 and the validated gRNAs targeting *Oga*. These cell lines ensure a timed knockout of *Oga* in a controllable manner (*Oga*-KO, fig. S2, A and B). Depletion of *Oga* led to enhanced O-GlcNAcylation in OPCs (fig. S2, C and D). As a result, the O4^+^ or Mbp^+^ cells were increased to 30.9±1.0 % and 20.4±1.6 % (21.2±0.6 % and 13.5±1.6 %, respectively, in vector control) (fig. S2, E and F). In addition, the resulting OLs exhibited mature morphology, with the branch number increased by 1.6-fold and the average length of processes increased by 1.8-fold (fig. S2, E, G and H). Furthermore, the expression of mature OL-markers were greatly induced, evidenced by imaging-based analysis (fig. S2, E and F) and qRT-PCR (4.2-fold for *Mbp*, 3.0-fold for *Mag*, and 3.2-fold for *Mog*) (fig. S2I).

**Fig. 2.**
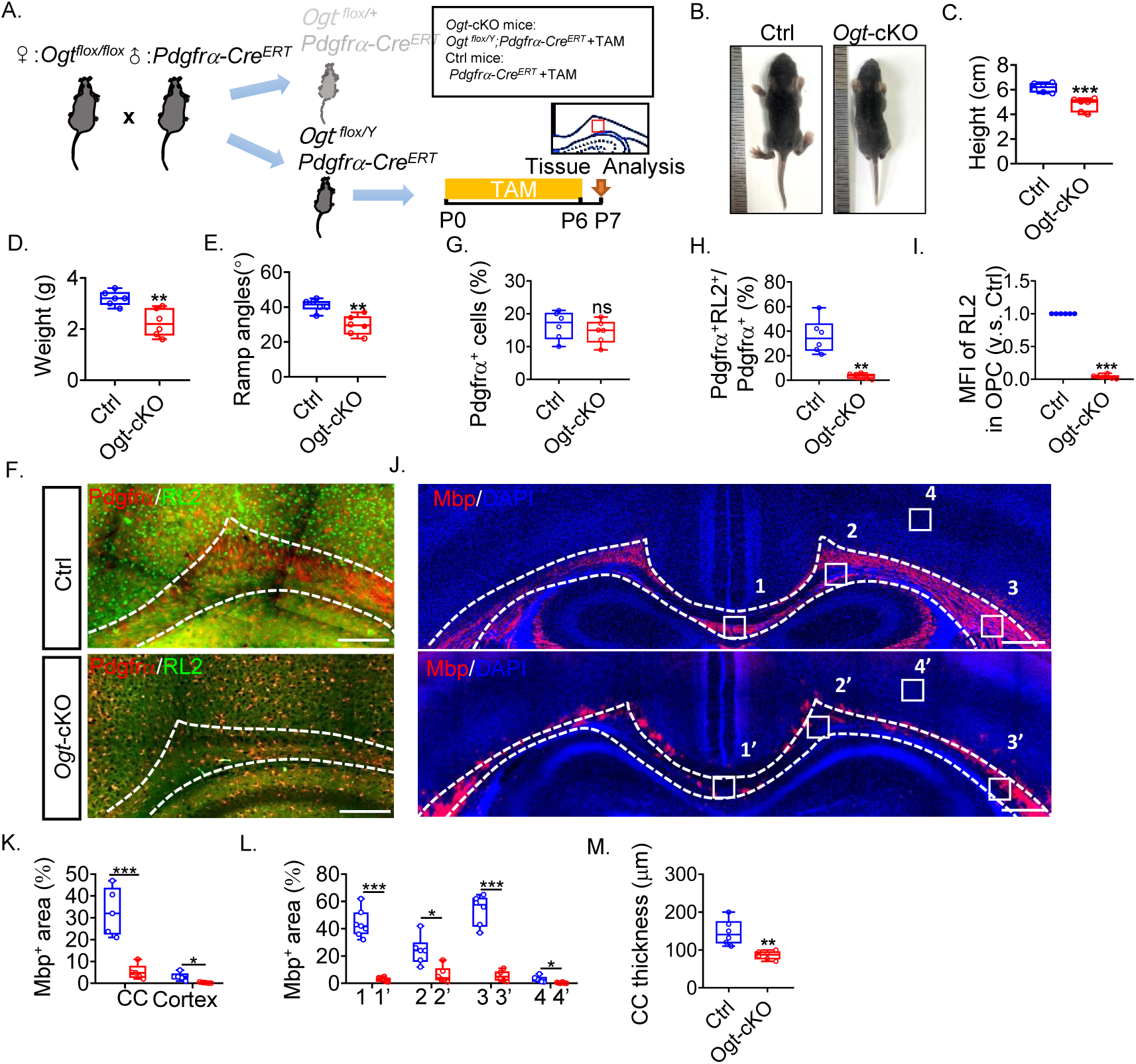
Ablation of *Ogt* in OPCs Impairs OPC Development and Myelin Development. (**A**) Schematic diagram shows the procedure for the generation of *Ogt*-cKO mice. Tamoxifen (TAM) was administered from P0 to P6 daily, and the brain and spinal cord (SC) tissues were collected and analyzed by P7. (**B-D**) Representative images of control (Ctrl, *Pdgfrα-Cre^ERT^*+TAM) and *Ogt*-cKO mice (*Ogt ^flox/Y^;Pdgfr*α*-Cre^ERT^*+TAM) (**B**) and the quantification of their height (**C**) and weight (**D**). n=6 mice per group. (**E**) The average maximum ramp angles from the inclined plane test for Ctrl and *Ogt*-cKO mice were quantified, respectively. n=6 mice per group. (**F-I**) Immunofluorescence analysis of OPCs and O-GlcNAcylated proteins (**F**) from the mouse brains of Ctrl or *Ogt*-cKO mice, respectively. The dashed lines in the left panel indicate the corpus callosum (CC). Scale bars, 100 μm (**F**). The percentage of Pdgfrα^+^ cells (**G**), Pdgfrα^+^RL2^+^/Pdgfrα^+^ cells (**H**) in the CC, and the fold enrichment of MFI of RL2 in OPCs compared to that in Ctrl group (**I**) were quantified. n=6 mice per group. Five fields of confocal images per sample were analyzed. (**J-M**) Immunofluorescence analysis of Mbp^+^ cells in the CC and cortex of Ctrl or *Ogt-*cKO mice (**J**). The dashed lines indicated the CC. Scale bars, 500 μm. The percentage for Mbp^+^ areas in the whole area in (**J**) of CC and Cortex were quantified in (**K** and **L**), respectively. The mean thickness of CC was quantified in (**M**). n=6 mice per group. All the quantification data are presented as mean ± SEM. *p*-values were calculated using two-tailed unpaired Student’s t-test (**C-E, G-I, K-M**). \**p* < 0.05, \*\**p* < 0.01, \*\*\**p* < 0.001, ns: not significant.

Meanwhile, we analyzed the differentiation of OPCs with genetic depletion of *Ogt* (*Ogt*-KO, fig. S2, A and J). Remarkably, knockout of *Ogt* reduced the O-GlcNAcylation globally (fig. S2, C and D) and seriously impeded OPC differentiation (O4^+^ and Mbp^+^ cells were reduced to 1.8 % and 1.2 %, respectively) (fig. S2, E and F). The resulting OLs were with fewer and shorter processes on average (the number and average length of processes reduced to 22.2 % and 64.8 % of vector control, respectively) (fig. S2, E, G and H) and reduced expression of OL-specific genes (16 % for *Mbp*, 12 % for *Mag*, and 15 % for *Mog* of vector control, respectively) (fig. S2I). No significant difference in total cell number was observed in cultures between control and *Ogt*-KO groups, excluding the possibility that reduced differentiation efficiency in *Ogt*-KO OPCs was due to cell death (fig. S2K). These data collectively support that O-GlcNAcylation promotes OPC differentiation *in vitro*.

### Depletion of *Ogt* in OPCs Compromises Myelin Development

Given its function in promoting OPC differentiation, we examined whether O-GlcNAcylation regulates myelin development *in vivo*. To this end, the *Ogt^flox/flox^* female mice were crossed with *Pdgfrα-Cre^ERT^* male mice. We chose *Ogt ^flox/Y^*; *Pdgfrα-Cre^ERT^* offsprings for further study and used the *Pdgfrα-Cre^ERT^* mice as control (Fig. 2A). By P7, we observed retarded body weight/height and brain development in *Ogt*-cKO mice (Fig. 2, B to D and fig. S3A), which exhibited abnormality in locomotion through an inclined plane test (Fig. 2E). We did not observe significant changes in the number of OPCs at corpus callosum (CC), but a dramatic decrease of O-GlcNAcylation (Fig. 2, F-I), further supported a successful and specific depletion of *Ogt* in OPCs.

To analyze the myelin development, we evaluated the myelin basic protein (Mbp) at both CC and cortex. As we observed a severe attenuation of the myelin structure in the absence of *Ogt* (Fig. 2, J-M), the Mbp^+^ area reduced from 32.9 % to 5.2 %, and 3.1 % to 0.3 %, for CC and cortex, respectively (Fig. 2, J and K). Anatomically, CC in *Ogt*-cKO mice shrunk and the average thickness was reduced, which was confirmed by immunofluorescence (IF) (Fig. 2, J and M) and myelin staining (fig. S3, C and F), collectively suggesting impaired myelin formation. Moreover, the expression of myelin-related genes was dramatically decreased in the cortex, CC, and spinal cord, further demonstrating a severe impairment of myelination throughout the CNS (fig. S3B). This battery of results collectively demonstrated an essential role of O-GlcNAcylation in myelin development in the CNS.

### O-GlcNAcylation in OPCs Displays a Positive Correlation with Myelin Status

To further analyze the function, we assessed the correlation of O-GlcNAcylation with myelin status. We applied a toxin-induced model by injecting lysolecithin (LPC) into the CC of the mouse brain (LPC mice, Fig. 3, A and B). Injection of LPC led to instant demyelination in CC (marked by Iba1 staining). OPCs repopulated and initiated to differentiate in the lesions around 10 days post-lesion (d.p.l.), and remyelination was evident by ∼ 14 d.p.l. and nearly completed by ∼ 21 d.p.l. (Fig. 3, C and D). We would, therefore, be able to monitor the dynamics of O-GlcNAcylation in OPCs over two stages — OPC repopulation (from 0 to 10 d.p.l.) and differentiation (from 10 to 21 d.p.l.). OPCs increased in the lesion at the early stage (0 to 10 d.p.l.), peaking at ∼ 4 to 7 d.p.l., indicating a repopulation (Fig. 3E). Among those cells, however, the density and percentage of RL2^+^ OPCs dropped significantly (Fig. 3, F and G). Meanwhile, the level of O-GlcNAcylation (mean fluorescence intensity, MFI) in those RL2^+^ OPCs was decreased, concomitantly with the myelin loss (Fig. 3H). From ∼ 10 d.p.l. on, active remyelination was initiated, evidenced by the reduced area of the demyelination (Fig. 3, C and D). Notably, although the number of OPCs did not change significantly, the density and percentage of RL2^+^ OPCs in lesions increased gradually (Fig. 3, F and G). Until 21 d.p.l., the RL2^+^ OPCs became comparable to those in WT mice (Fig. 3, F and G). Furthermore, O-GlcNAcylation in those OPCs was upregulated along with remyelination (Fig. 3H). Together, these results revealed that O-GlcNAcylation in OPCs shows a positive correlation with myelin status.

**Fig. 3.**
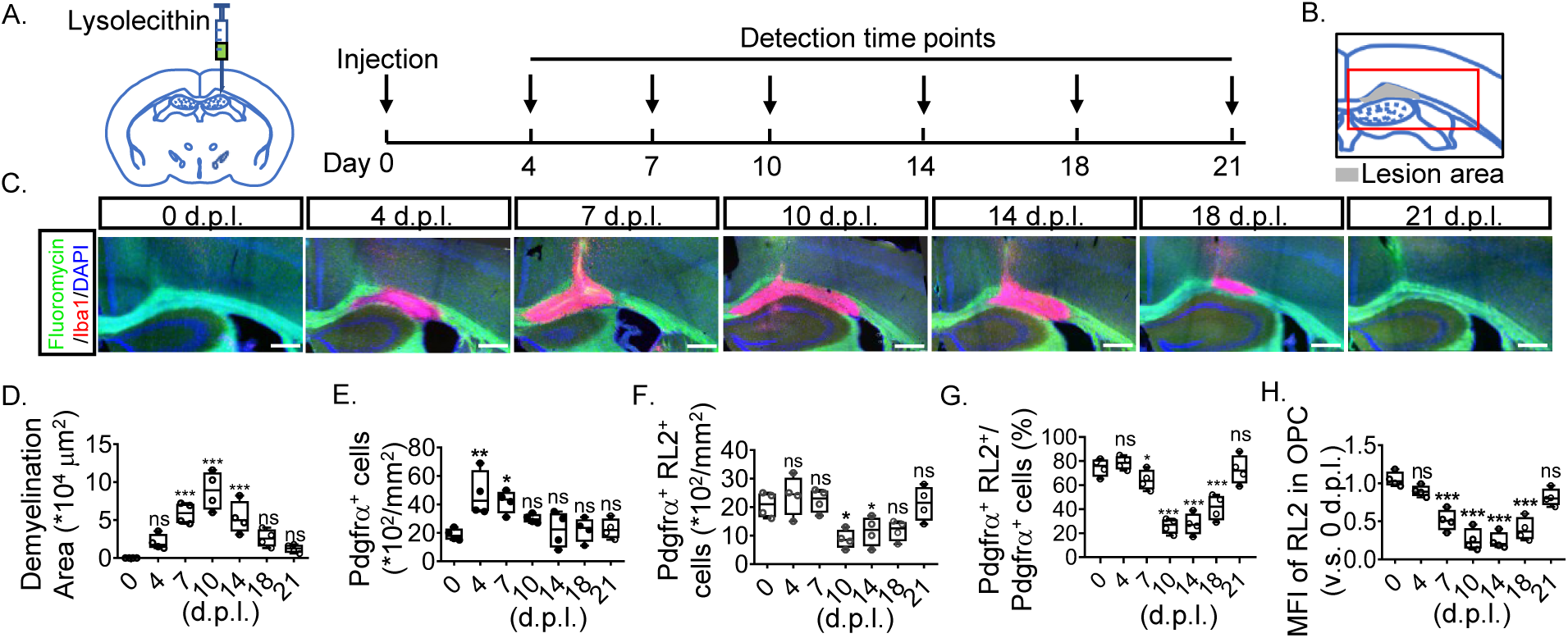
O-GlcNAcylation Is Positively Correlated with Myelin Status. (**A, B**) Time course analysis of the O-GlcNAcylation in OPCs in the LPC-induced mice model. The lesion at CC that was defined by Iba1 staining was analyzed by 0, 4, 7, 10, 14, 18, and 21 d.p.l. (**A**). The demyelination region was marked in gray, and the imaged region was marked in red in (**B**). (**C, D**) Myelin (Fluoromycin) and lesion (Iba1^+^ cells) in CC were analyzed through immunofluorescence analysis (**C**) and the areas of demyelination regions were quantified in (**D**). Four mice (two male mice and two female mice of eight weeks) were used for each time point. Scale bars, 200 μm in (**C**). (**E-H**) Immunofluorescence analysis of OPCs and O-GlcNAcylation in lesion at indicated time points. The density of total OPCs (**E**), RL2^+^ OPCs (**F**), the percentage of RL2^+^ OPCs (Pdgfrα^+^ RL2^+^/Pdgfrα^+^ cells) (**G**), the fold enrichment of MFI of RL2 in OPCs (**H**) respectively, in the lesion, were quantified. The statistical difference was evaluated by comparing it to those at 0 d.p.l. Four mice of eight weeks were used for each group. All the quantification data are presented as mean ± SEM. *p*-values were calculated using a one-way ANOVA test (**D-H**). \**p* < 0.05, \*\**p* < 0.01, \*\*\**p* < 0.001, ns: not significant.

### Upregulation of O-GlcNAcylation Drives Remyelination

Considering its positive correlation with myelin status, we assessed if O-GlcNAcylation in OPCs would regulate remyelination. To this end, we first applied the small-molecule modulators Thiamet-G (TMG, an Oga inhibitor, 10 μM) or OSMI-1 (an Ogt inhibitor, 12.5 μM), respectively, to the OPC differentiation assay. ESC-derived OPCs were treated with TMG or OSMI-1 to mimic an enhanced or reduced O-GlcNAcylation, respectively. Treating cells with TMG dramatically promoted OPC differentiation (fig. S4, A to C), and the resulting OLs had an increased number and length of branches, resembling a more mature phenotype (fig. S4, A, D and E). This effect of TMG is dose-dependent, suggesting a specific induction (fig. S4, H, J and K). qRT-PCR revealed a dramatic increase in the expression of defining genes for mature OLs with TMG, supporting a robust differentiation (fig. S4G). In contrast, treating OPCs with OSMI-1 remarkably impaired the generation of OLs in a dose-dependent manner, and resulted in immature OLs with lower expression of myelin-related genes (fig. S4, A to E, G, I, L, M). Few mature OLs were detected even with extended differentiation, excluding the possibility that OSMI-1 slowed down the dynamics of differentiation (data not shown).

**Fig. 4.**
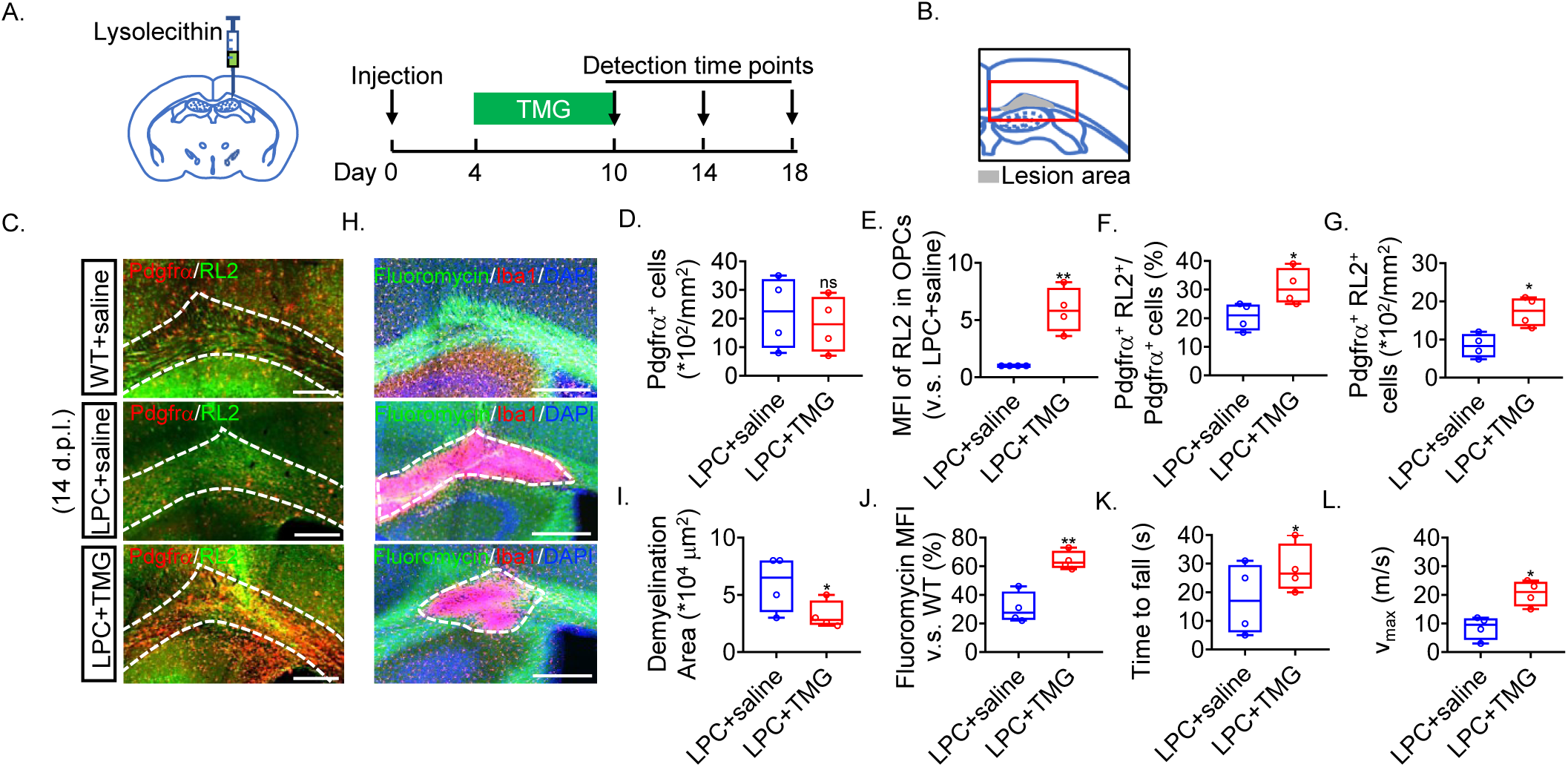
Pharmacological Upregulation of O-GlcNAcylation Drives Remyelination. (**A, B**) The diagram shows the procedure for validating the function of TMG on remyelination in the LPC-induced mice model (**A**). The demyelination region was marked in gray, and the imaged region was marked in red in (**B**). (**C-G**) Immunofluorescence analysis for OPCs and O-GlcNAcylated proteins (**C**) from the mouse brains of the WT+saline, LPC+saline, and LPC+TMG mice, respectively. Scale bars, 200 μm (**C**). The dashed lines in the panel indicate the CC. The density of Pdgfrα^+^ OPCs (**D**), normalized MFI of RL2 in Pdgfrα^+^ OPCs (**E**), percentage of RL2^+^ OPCs (Pdgfrα^+^RL2^+^/Pdgfrα^+^) (**F**), and the density of Pdgfrα^+^RL2^+^ cells (**G**) in the CC were quantified. Four mice of eight weeks were used for each group. (**H-J**) Myelin (Fluoromycin) in the lesion (Iba1^+^ cells) at the CC for control (WT+saline) or LPC-treated mice receiving either saline or TMG, respectively, were analyzed by immunofluorescence (**H**). The dashed lines indicate the demyelination region of CC for quantification. Scale bars, 200 μm. The area of demyelination regions (**I**) compared to the WT group was quantified. The normalized MFI of Fluoromycin in the lesion based on (**H**) was quantified (**J**). Four mice of eight weeks were used for each group. (**K, L**) The analyses of the rotating rod test for fixed speed (**K**) and accelerating test (**L**) were shown. Four mice of eight weeks were used for each group. All the quantification data are presented as mean ± SEM. *p*-values were calculated using two-tailed unpaired Student’s t-test (**D-G, I-L**). \**p* < 0.05, \*\**p* < 0.01, \*\*\**p* < 0.001, ns: not significant.

Next, we examined if pharmacological upregulation of O-GlcNAcylation could drive remyelination *in vivo*. In the LPC-induced demyelination mouse model, the lesioned mice sustained serious myelin disruption by 14 d.p.l. TMG (20 mg per kg body weight) was intraperitoneally injected daily from 4 to 10 d.p.l. (Fig. 4, A and B). By 14 d.p.l., although the number of OPCs did not change greatly, administration of TMG induced O-GlcNAcylation dramatically in OPCs (Fig. 4, C to E), and the percentage and density of RL2^+^ OPCs increased greatly (Fig. 4, F and G). Consequently, remarkable remyelination was observed with significantly reduced lesion area and increased myelin (Fig. 4, H to J). Meanwhile, the animals exhibited considerable recovery on locomotion (Fig. 4, K and L), suggesting a vital role of O-GlcNAcylation on remyelination. These batches of data provided compelling evidence that pharmacological upregulation of O-GlcNAcylation drives remyelination.

### O-GlcNAcylation Intrinsically Drives Remyelination

The pharmacological induction of O-GlcNAcylation by TMG may not be specific to OPCs. We, therefore, examined if genetic manipulation of O-GlcNAcylation in OPCs could modulate remyelination. We first assessed how remyelination proceeds in the absence of *Ogt*. We treated *Ogt ^flox/Y^*; *Pdgfrα-Cre^ERT^* mice (8 weeks old) with TAM (defined as day 0) to produce the *Ogt*-cKO adult mice (used *Pdgfrα-Cre^ERT^* mice as control). Demyelination was induced by LPC injection at day 35 in mice with (ctrl) or without *Ogt* (*Ogt*-cKO). These two groups of mice were then administered with saline or TMG, respectively, from day 39 for six consecutive days. The extent of remyelination at CC was analyzed by day 49 (Fig. 5A). In the presence of *Ogt* (Groups A and B), treating mice with TMG promoted remyelination (Fig. 5, B and C). In contrast, in the absence of *Ogt* (Groups C and D), TMG was not that effective, and severe demyelination was sustained, suggesting O-GlcNAcylation is necessary for remyelination (Fig. 5, B and C). Together, these results consistently demonstrated that O-GlcNAcylation in OPCs is necessary for remyelination.

**Fig. 5.**
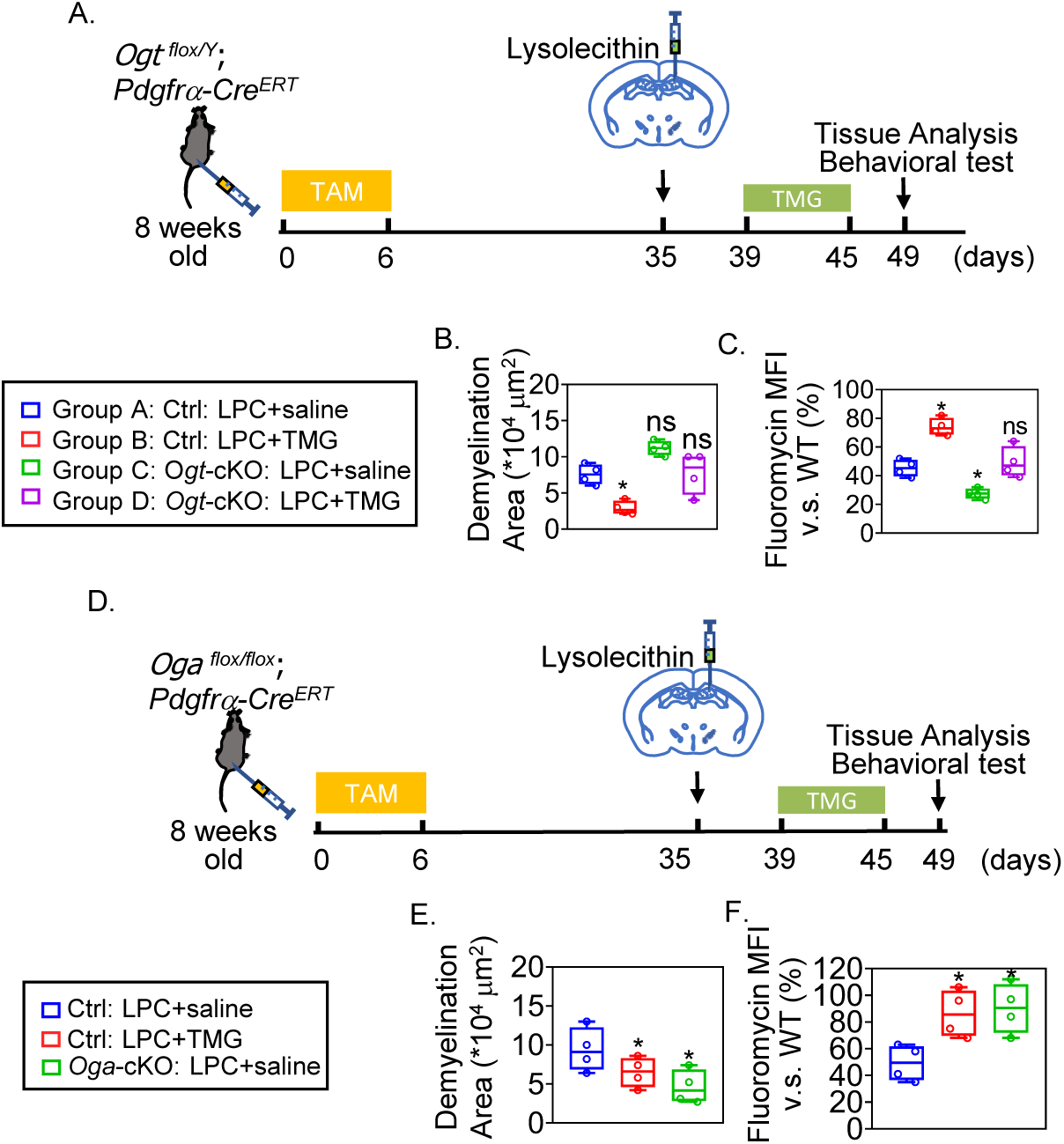
Ablation of *Ogt* in OPCs Compromises Remyelination. (**A**) The procedure of the analyses of the function of *Ogt* in the LPC model. (**B, C**) Myelin (Fluoromycin) in the lesion (Iba1^+^ cells) at the CC for indicated groups was analyzed by immunofluorescence. The area of demyelination regions (**B**) and the normalized MFI of myelin (Fluoromycin) (**C**) were quantified. Three mice of eight weeks were used for each group. (**D**) The procedure of the analyses of the function of *Oga* in the LPC model. (**E, F**) Myelin (Fluoromycin) in the lesion (Iba1^+^ cells) at the CC for indicated groups were analyzed by immunofluorescence. The area of demyelination regions (**E**) and the normalized MFI of Fluoromycin (**F**) were quantified. Four mice of eight weeks were used for each group. Four mice of eight weeks were used for each group. All the quantification data are presented as mean ± SEM. *p*-values were calculated using one-way ANOVA test (**B, C, E, F**). \**p* < 0.05, \*\**p* < 0.01, \*\*\**p* < 0.001, ns: not significant.

Next, we explored if the upregulation of O-GlcNAcylation in OPCs is sufficient to drive remyelination. To this end, we generated *Oga*-cKO adult mice by crossing *Oga^flox/flox^* with *Pdgfrα-Cre^ERT^* mice (Fig. 5D). These *Oga*-cKO adult mice were injected with LPC at CC at day 35 to induce demyelination, then administered daily with TMG or saline from day 39 to 45, and the extent of remyelination were assessed by day 49 (Fig. 5D). By day 49, as the demyelination lesion recovered more efficiently in *Oga*-cKO adult mice than in the control mice receiving either saline or TMG, the myelin fibers were more evident in the lesion as well (Fig. 5, E and F). Together, these genetic data demonstrate that the upregulation of O-GlcNAcylation in OPCs is sufficient to drive remyelination.

### Tet3 Is Verified as One O-GlcNAcylated Protein Mediating Myelin Formation

To identify the factors in OPCs that mediate the O-GlcNAcylation-induced remyelination, we employed an O-GlcNAcylation-specific proteomics analysis. The total proteins from purified OPCs of P0 mice were trypsinized, and the O-GlcNAcylated peptides were enriched using O-GlcNAcylation antibodies conjugated beads and subjected to liquid chromatography with tandem mass spectrometry (LC-MS/MS) analysis. We identified Tet3 as one O-GlcNAcylated protein and further verified that the knockout of *Tet3* impaired the generation of OLs significantly, supporting their involvement in OPC differentiation. We, therefore, explored the function of Tet3.

To explore whether O-GlcNAcylation of Tet3 affects OPC differentiation, we isolated the primary NPCs (pNPCs) from the brains of *Tet3^flox/flox^* fetal mice at E13.5. The plasmids encoding Cre recombinase and either Tet3^WT^ or Tet3 mutant (the O-GlcNAcylated Serine was mutated to Alanine, Tet3^S-A^), respectively, were transfected into the pNPC-derived OPCs. Therefore, the endogenous *Tet3* could be disrupted in OPCs (OPC^Tet3-KO^), and the effects of Tet3 (OPC^Tet3-WT^) or its mutation (OPC^Tet3-S-A^) on OPC differentiation could be assessed. As a result, OPC^Tet3-KO^ exhibited a dramatically compromised capacity of differentiation (O4^+^ or Mbp^+^ cells were 1.6 % and 0.9 %), supporting the important role of Tet3 in OPC differentiation. Meanwhile, the differentiation efficiency was significantly reduced for OPC^Tet3-S-A^ (O4^+^ or Mbp^+^ cells were 3.7 % and 3.0 %, respectively, which were 20.3 % and 14.5 %, respectively, for OPC^Tet3-WT^) (Fig. 6, A to C), suggesting that O-GlcNAcylation of Tet3 is crucial for its function on OPC differentiation. Of note, unlike OPC^Tet3-WT^, treating OPC^Tet3-S-A^ with TMG exhibited little effect on promoting differentiation, providing compelling evidence that O-GlcNAcylated Tet3 is crucial for O-GlcNAcylation-induced myelin formation.

**Fig. 6.**
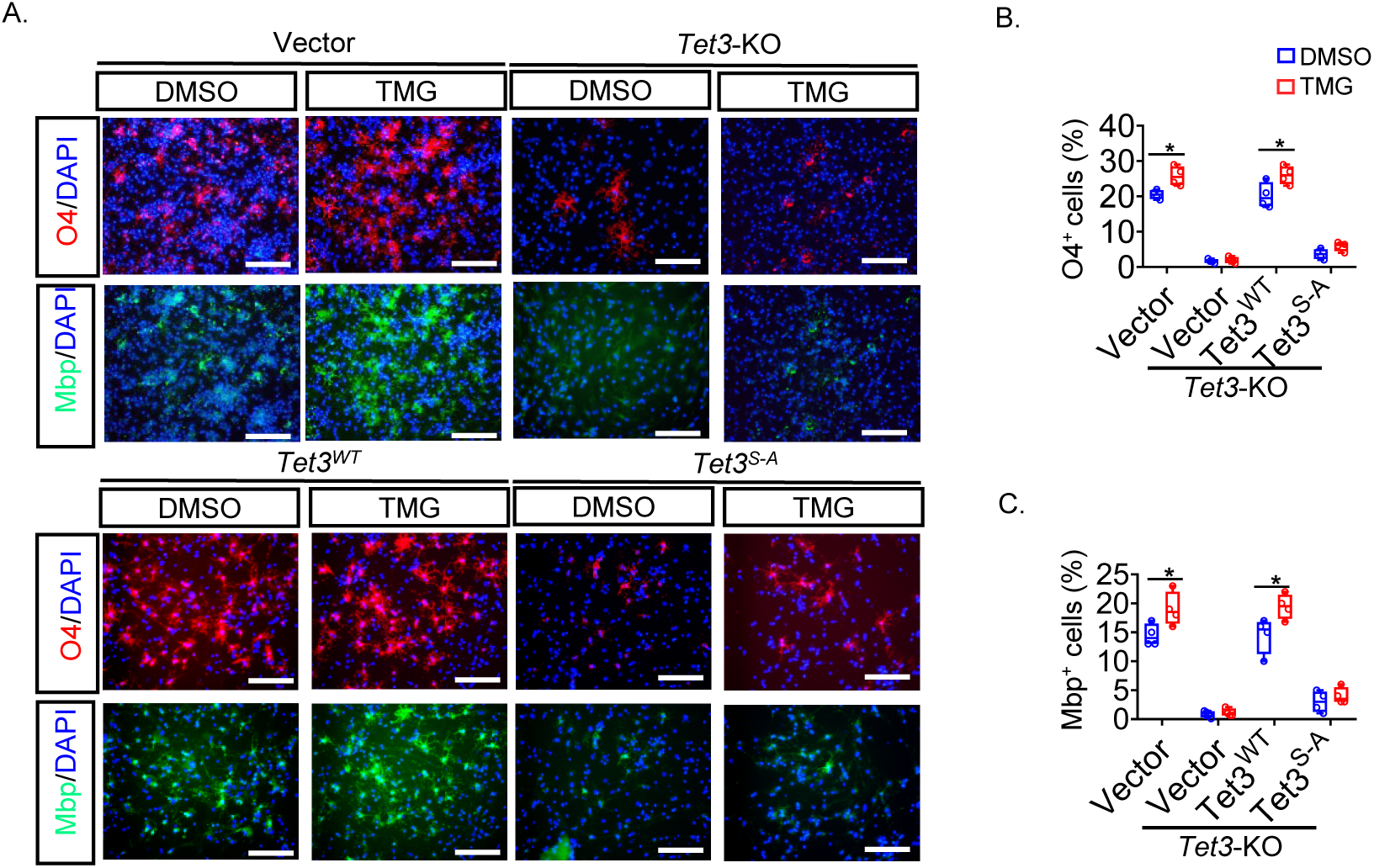
Tet3 Is One O-GlcNAcylated Factor Inducing OPC Differentiation. (**A-C**) Representative images show the differentiation efficiencies of OPCs under indicated conditions (**A**). Scale bars, 50 μm. The percentage of O4^+^ (**B**) or Mbp^+^ (**C**) cells was quantified. n= 4 independent experiments. All the quantification data are presented as mean ± SEM. *p*-values were calculated using two-tailed unpaired Student’s t-test (**B, C**). \**p* < 0.05, \*\**p* < 0.01, \*\*\**p* < 0.001, ns: not significant.

### O-GlcNAcylation of Tet3 Safeguards Its Catalytic Activity and Ensures the Myelin Gene Expression

To clarify the mechanism of Tet3 in regulating myelin gene expression, we investigated if O-GlcNAcylation affects the catalytic activity of Tet3. We transduced OPC^Tet3-KO^ with Flag-tagged *Tet3*^WT^ or *Tet3*^S-A^. The genomic DNA from both OPC^Tet3-KO^ and OPC^Tet3-S-A^, respectively, were isolated for DNA dot blotting analysis with antibodies against 5hmC (Fig. 7A). Consistently, we found that OPC^Tet3-S-A^ showed a reduced level of 5hmC, supporting that loss of O-GlcNAcylation leads to impaired catalytic activity of Tet3. On the other hand, to elucidate whether Tet3^S-A^ would lead to an increased level of 5-methylcytosine (5mC), we performed a DNA bisulfite sequencing analysis for the promoter of myelin genes. Accordingly, we detected a higher deposition of 5mC at the promoter of the myelin genes, such as *Sox10*, *Pdgfrα*, and *Olig2*, than that in OPC^Tet3-WT^ (Fig. 7B). As a consequence, the expression of the myelin genes was dramatically repressed in OPC^Tet3-S-A^. Collectively, our finding establishes a crucial role of the Ogt-Tet3 axis in driving remyelination, where O-GlcNAcylation of Tet3, one downstream factor of Ogt, safeguards OPC differentiation, at least in part, through modulating its catalytic activity and consequently regulates OPC differentiation and remyelination.

**Fig. 7.**
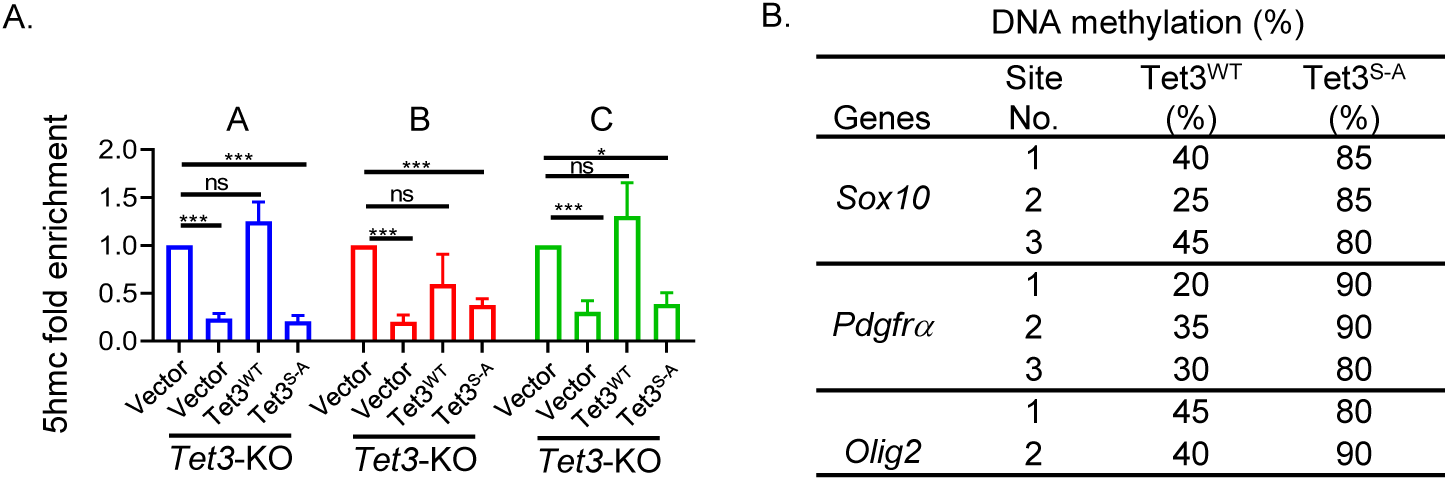
O-GlcNAcylation of Tet3 Safeguards Its Catalytic Activity and OPC Differentiation. (**A**) DNA dot blotting analyses show the level of 5hmC under indicated conditions. n = 3 independent samples. (**B**) The bisulfate sequencing analyses show the methylation of promoter regions for indicated conditions. Open circle, nonmethylated CpG; closed circle, methylated CpG. n = 20 replicates for each site. All the quantification data are presented as mean ± SEM. *p*-values were calculated using one-way ANOVA test (**A**), \**p* < 0.05, \*\**p* < 0.01, \*\*\**p* < 0.001, ns: not significant.

---

**Supplementary Fig. 1.**
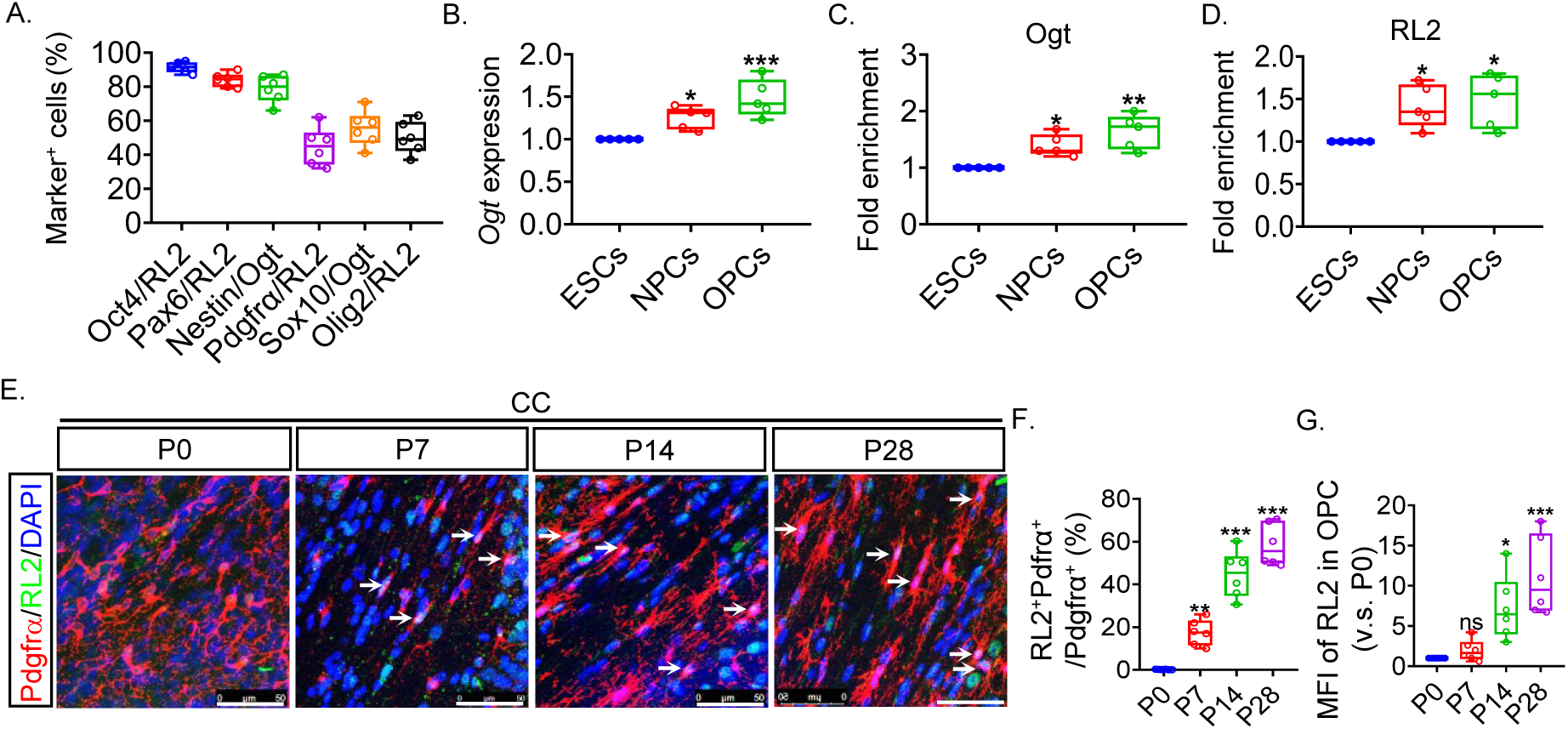
O-GlcNAcylation Modification for Proteins in OPCs. Related to. **Fig. 1**. (**A**) The percentage of cells co-expressing indicated cell marker and RL2 or Ogt was quantified based on immunostaining results. n= 6 independent biological experiments. (**B**) qRT-PCR analysis for the expression of *Ogt* in the indicated cell types. The relative expression was normalized to that in ESCs. n=5 independent experiments. (**C, D**) Quantification of the expression of Ogt (**C**) or O-GlcNAcylated proteins (**D**) in indicated cell types based on Western blot results. n=5 independent experiments. (**E-G**) Immunofluorescence images show the O-GlcNAcylation in Pdgfrα-expressing OPCs from the corpus callosum (CC) of mice at P0, P7, P14, and P28 (**E**), respectively. The percentage of RL2^+^Pdgfrα^+^/Pdgfrα^+^ cells based on immunofluorescence results was quantified (**F**). The fold enrichment of MFI of RL2 in Pdgfrα^+^ OPCs based on immunofluorescence results compared to P0 was quantified (**G**). Scale bars=50 μm in (**E**). Six mice were used for each group. All the quantification data are presented as mean ± SEM. *p*-values were calculated using a one-way ANOVA with Tukey’s multiple comparisons test (**B-D, F, G**). \**p* < 0.05, \*\**p* < 0.01, \*\*\**p* < 0.001.

**Supplementary Fig. 2.**
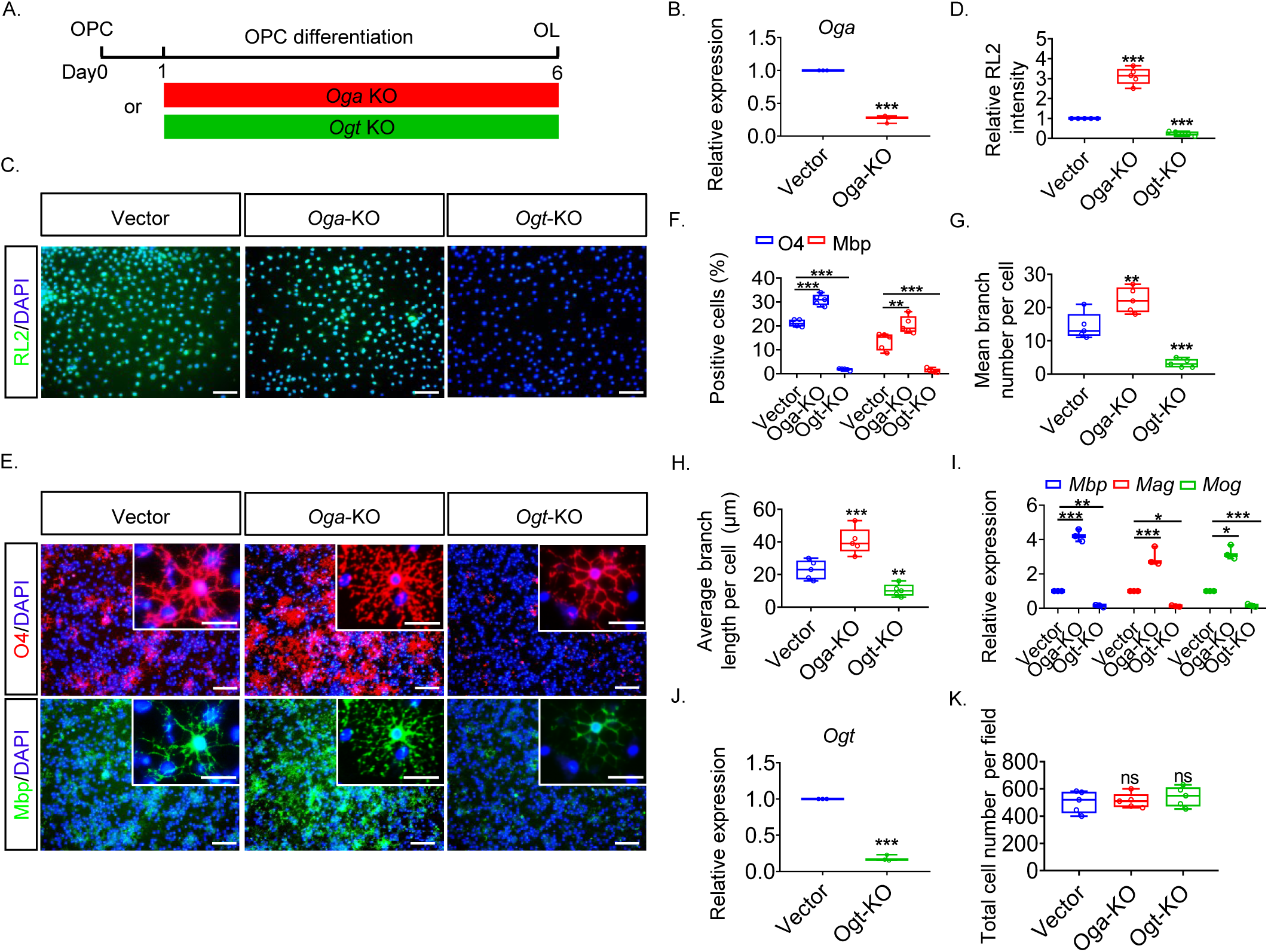
O-GlcNAcylation Promotes OPC Differentiation *in vitro*. (**A**) Schematic diagram for the evaluation of OPC differentiation with genetic depletion of *Oga* (*Oga*-KO) or *Ogt* (*Ogt*-KO), respectively. (**B**) The expression of *Oga* in OPCs under indicated conditions was analyzed by qRT-PCR and normalized to vector control. n=3 independent experiments. (**C, D**) Representative immunostaining results show the O-GlcNAcylation in OPCs with genetic depletion of *Ogt* or *Oga*, respectively (**C**). The relative intensity of RL2 was quantified and normalized to vector control (**D**). Scale bars, 50 μm in (**C**). n=5 independent biological experiments. (**E-H**) Representative immunostaining results show the OPC differentiation under indicated conditions (**E**). Scale bars, 50 μm and 25 μm for zoomed-in images in (**E**). The percentage of O4^+^ or Mbp^+^ cells (**F**), the number of branches per cell (**G**), and the average length of branches (**H**) were quantified. n=5 independent biological experiments. (**I**) The expression of indicated genes was analyzed by qRT-PCR and normalized to the vector control. n=3 independent experiments. (**J**) The expression of *Ogt* in OPCs under indicated conditions was analyzed by qRT-PCR and normalized to vector control. n=3 independent experiments. (**K**) The total cell number for indicated conditions was quantified. n=5 independent biological experiments. All the quantification data are presented as mean ± SEM. *p*-values were calculated using two-tailed unpaired Student’s t-test (**B, J**) and one-way ANOVA test (**D, F-I, K**). \**p* < 0.05, \*\**p* < 0.01, \*\*\**p* < 0.001, ns: not significant.

**Supplementary Fig. 3.**
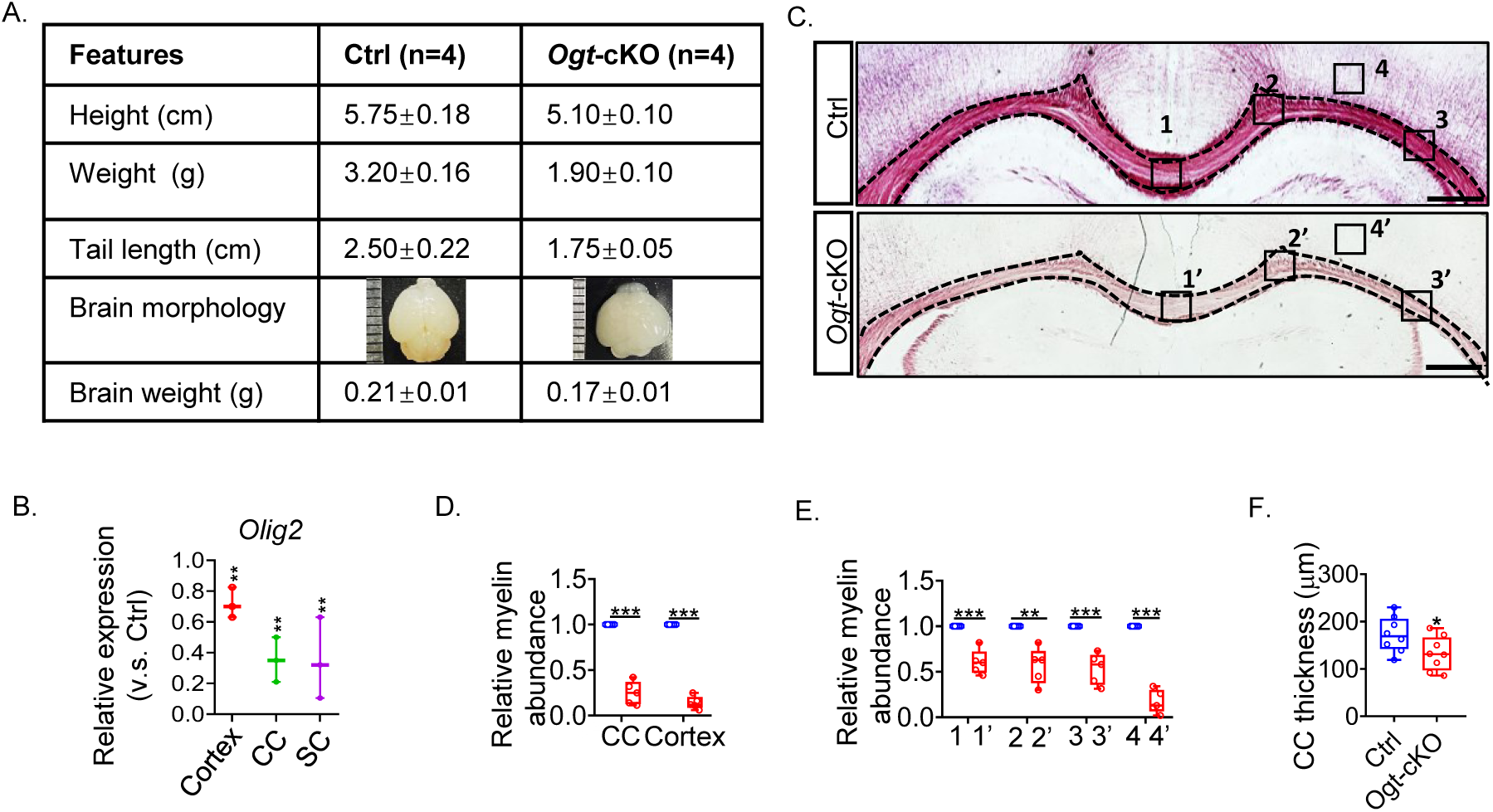
Ablation of *Ogt* in OPCs Impairs Myelin Development. Related to. **Fig. 2**. (**A**) Table describes the indicated information for control mice (Ctrl, *Pdgfrα-Cre^ERT^*) and *Ogt*-cKO mice. n=4 mice for each group. (**B**) The expression of indicated genes in samples from the cortex, CC, and SC of *Ogt*-cKO mice was analyzed by qRT-PCR and normalized to those of the Ctrl group. n=3 independent biological experiments. (**C-F**) Gold myelin staining analysis shows the myelin of Ctrl mice and *Ogt*-cKO mice at the CC and Cotex (**C**). Scale bars, 200 μm. The relative myelin abundance in (**C**) were shown in (**D**, **E**). The thickness of the CC in (**C**) was quantified in (**F**). n= 5 mice for each group. All the quantification data are presented as mean ± SEM. *p*-values were calculated using two-tailed unpaired Student’s t-test (**B, D-F**). \**p* < 0.05, \*\**p* < 0.01, \*\*\**p* < 0.001, ns: not significant.

**Supplementary Fig. 4.**
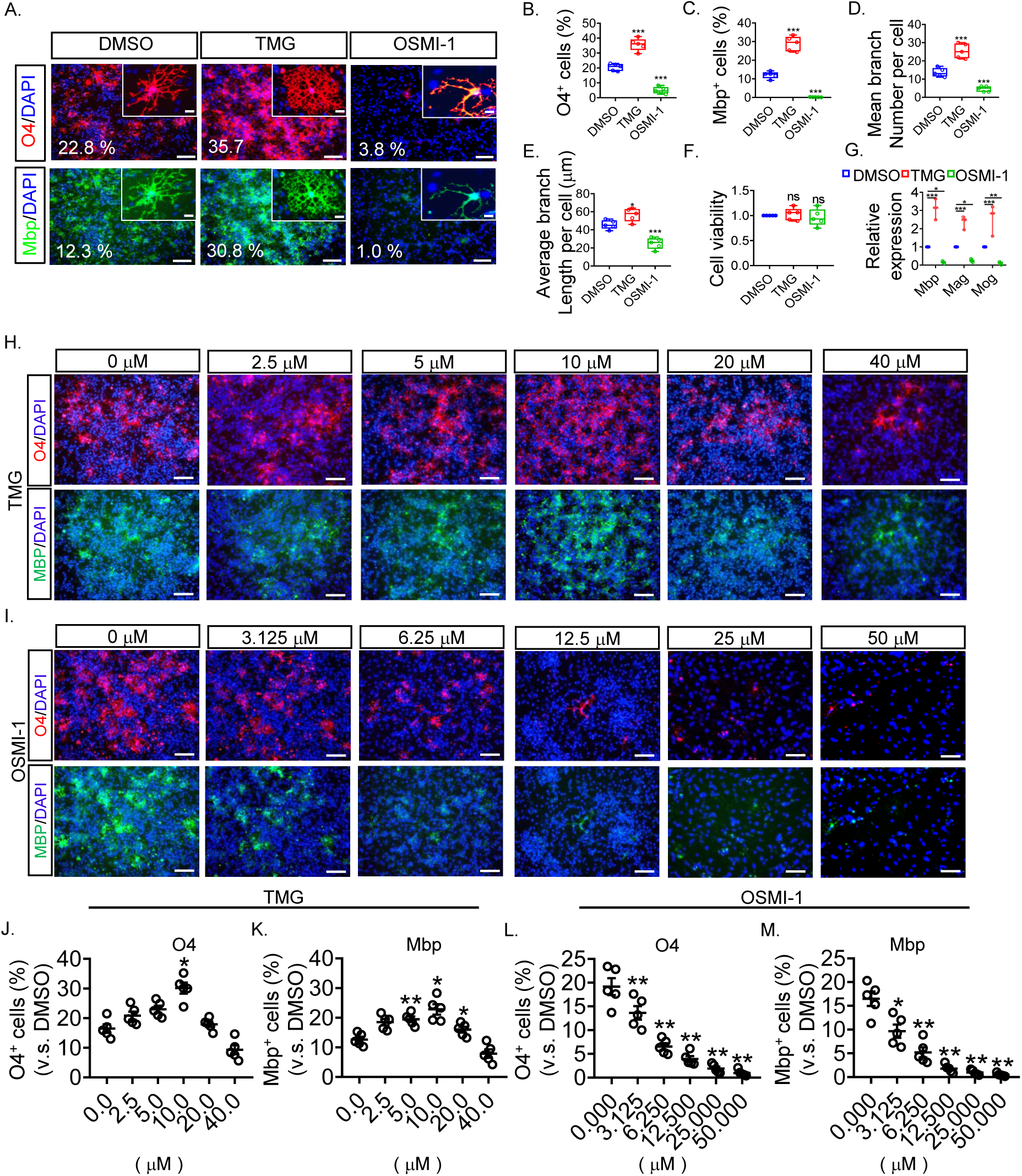
Effects of TMG and OSMI-1 on OPC Differentiation. (**A-E**) The immunostaining results show the effect of TMG and OSMI-1 on OPC differentiation. Scale bars, 50 μm, and 25 μm in zoomed-in images in (**A**). The percentage of O4^+^ or Mbp^+^ cells under indicated conditions is shown in Figures. The percentages of O4^+^ cells (**B**), Mbp^+^ cells (**C**), the mean branch number per cell (**D**), and the average length of branches (**E**) were quantified. n=5 independent biological experiments. (**F**) The cell viability was quantified and normalized to the DMSO group. n = 5 independent samples. (**G**) The expression of OL genes under indicated conditions was analyzed by qRT-PCR and normalized to the DMSO control group. n=3 independent experiments. (**H-M**) Immunostaining results show the dose effect of TMG (**h**) or OSMI-1 (**I**), respectively, on OPC differentiation. Scale bars, 50 μm. The percentage of O4^+^ (**J**, **L**) or Mbp^+^ (**K**, **M**) cells treated with TMG (**J, K**) or OSMI-1 (**L, M**), respectively, were quantified and normalized to the DMSO control group (0 μM). n= 5 independent biological experiments. All the quantification data are presented as mean ± SEM. *p*-values were calculated using a one-way ANOVA test (**B-G**, **J-M**). \**p* < 0.05, \*\**p* < 0.01, \*\*\**p* < 0.001, ns: not significant.

